# Role of interaction network topology in controlling microbial population in consortia

**DOI:** 10.1101/288142

**Authors:** Xinying Ren, Richard M. Murray

## Abstract

Engineering microbial consortia is an important new frontier for synthetic biology given its efficiency in performing complex tasks and endurance to environmental uncertainty. Most synthetic circuits regulate populational behaviors via cell-to-cell interactions, which are affected by spatially heterogeneous environments. Therefore, it is important to understand the limits on controlling system dynamics and provide a control strategy for engineering consortia under spatial structures. Here, we build a network model for a fractional population control circuit in two-strain consortia, and characterize the cell-to-cell interaction network by topological properties, such as symmetry, locality and connectivity. Using linear network control theory, we relate the network topology to system output’s tracking performance. We analytically and numerically demonstrate that the minimum network control cost for good tracking depends on locality difference between two cell population’s spatial distributions and how strongly the controller node contributes to interaction strength. To realize a robust consortia, we can manipulate the environment to form a strongly connected network. Our results ground the expected cell population dynamics in its spatially organized interaction network, and inspire directions in cooperative control in microbial consortia.

## I. Introduction

Synthetic biology provides biotechnologies to engineer cells and organisms with pre-specified functions. Interest has recently emerged in designing synthetic circuits for collective behaviors in consortia, which include metabolites trading for mutualism in co-culture [1], programmed predation and rescue in prey-predator systems [2], and other competition or coordination in sender-receiver systems [3]. Multiple populations communicate with each other, divide labor and perform subtasks so that the overall control goal can be achieved in consortia. Therefore, engineering microbial consortia will enrich the potential of synthetic gene circuits in gene therapy, biofuel and other applications [4], [5].

Realizing proposed behaviors of multiple interacting strains on diverse temporal or spatial scales relies on two perspectives. First, synthetic gene circuits need to be engineered in each population to exhibit certain functions. Second, it is essential to engineer cell-to-cell interaction to construct consortia. Bacteria have evolved interactions by producing, exporting and sensing autoinducers that are mostly small and diffusible signaling molecules. This phenomenon is called quorum sensing, where the accumulation of autoinducers in environment provides a good measure of cell population density and can quantitatively affect a cell’s gene expression or trigger a differentiation process [6]. Existing work in ecology demonstrate that the cooperative behaviors in consortia rely on the spatial distribution of multiple cell agents and the structure of their interaction [7]. Efforts in network science provide a good approach to define this spatial cell-to-cell interaction as a network graph, to describe the consortia as an integrated system and to analyze population level performance and robustness by graph metrics [8]. As multiple cell populations are usually spatially organized and form heterogeneous consortia, understanding the relationship between cell-to-cell interaction network topology and cooperative dynamics in consortia is particular useful for system-level controller design in synthetic biology. By linking the interaction network topology to populational behaviors, one can optimize control objectives by rearranging the spatial distribution of cell agents, rewiring intercellular quorum sensing pathways, reweighting interaction signaling strengths and turning on or off the synthetic circuits in certain cell agents via external chemical stimulations.

Recent research on engineering microbial consortia mainly focuses on intracellular circuit design [9], [10]. These papers assume that cell populations are well-mixed, because it is elusive how structured network properties enhance control potentials in spatial-organized consortia. Here, we propose a network control strategy of existing synthetic circuits and take different spatial distributions of cell agents and their interaction structures into consideration. Although the principles derived here are based on a specific synthetic circuit that regulates cell population via toxin/antitoxin mechanism, they can be applied to a more general design framework, where the autoinducers in quorum sensing and their regulated products are included in a feedback controller at DNA, transcriptional and translational levels.

This paper is organized as follow. In Section II, we give an overview of the biological design of the synthetic circuit. It performs as a fractional population controller in *E.coli* [11], [12]. In Section III, we construct a reaction-diffusion model under spatial settings to describe the full dynamics of the system. In Section IV, we derive a reduced network model and define network topology properties such as symmetry, locality and connectivity. In Section V, we use linear network control theory to obtain a mathematical approximation of the closed-form solution for system steady state. Our results include a computational simulation and a mathematical proof relating the interaction network topology properties to steady state cell populations. The results also include a graph representation of network topologies that are robust to perturbations. We summarize the main findings of the paper discuss future work in Section VI.

## II. Biological Design

To engineer a stable co-culture of two cell strains *Cell*_1_ and *Cell*_2_ characterized by referenced fractional populations, we require a synthetic circuit that consists of a biological sensor, a population comparator/controller and a cell growth actuator. The sensors are achieved by identifying a small library of quorum sensing signaling molecules (acyl homoserine lactones, AHLs) and corresponding receptor-promotor pairs. Two cell strains produces orthogonal AHLs respectively and concentrations of AHLs are approximations for their population densities. Both AHLs can bind with constitutive promoters to activate expression of toxin ccdB and antitoxin ccdA [13]. The comparator/controller is constructed by sequestration of the toxin/antitoxin pair to compare two scaled up cell population densities. The toxin ccdB functions as the actuator to regulate cell growth by inhibiting cell proliferation. The synthetic circuit is shown in Fig. 1(a). The reference is set by the relative level of external chemical inducer IPTG that induces two AHL synthesis. The circuit has good tracking accuracy of the reference and robust adaptation because it includes a sequestration-based controller that can be approximated as an integral feedback [14]–[16].

**Fig. 1.**
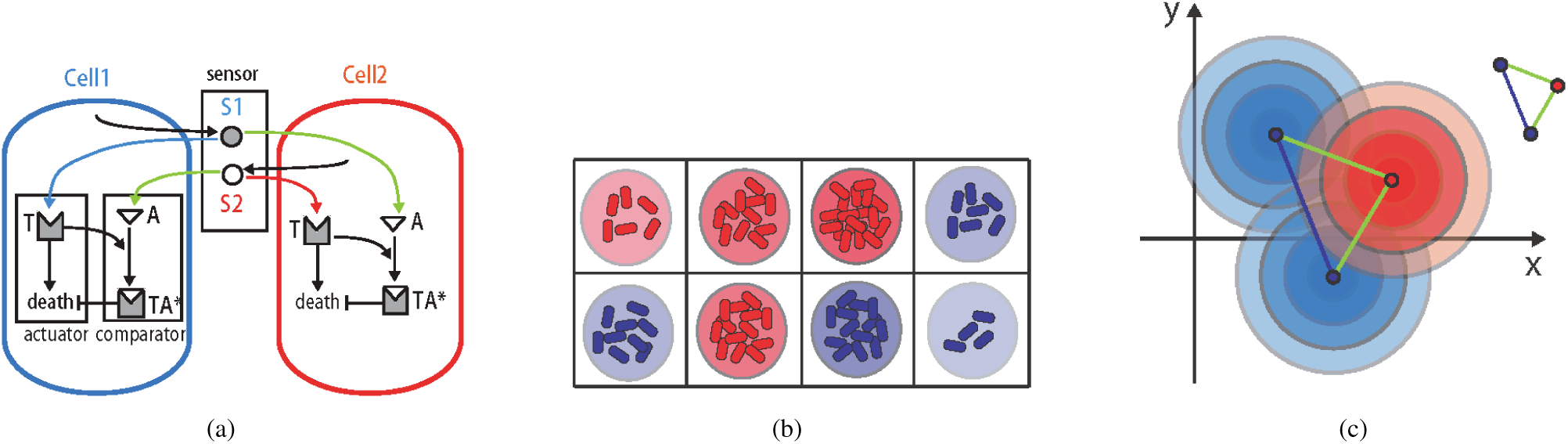
Abstract biological design of fractional population control circuit and spatial settings. (a) describes toxin-antitoxin sequestration based fractional population control circuit. Two cell population communicate via two quorum sensing signals *S*_1_,*S*_2_. In *Cell*_1_, *S*_1_ activates toxin *T* and *S*_2_ activates antitoxin A production to further regulate cell death and rescue processes through *T* and *A* sequestration. Similar reactions occur in *Cell*_2_. (b) describes the spatial setting of microcolonies cultured on an ager plate. Two cell populations are grown in separate wells where quorum sensing signals can diffuse around and accumulate in environment to form a certain concentration distribution on 2D space. (c) describes the diffusion of AHL molecules and the color shades represent AHL concentration levels. Cell microcolonies are represented as point sources of AHLs and the concentration of AHLs decreases along distance to the source following the diffusion law. The colored lines between microcolonies represent inter-strain or intra-strain interactions.

## III. Reaction-Diffusionmodel

We consider *N* microcolonies of strain *Cell*_1_ and *N* microcolonies of strain *Cell*_2_ cultured within 2*N* wells on an agar plate, as in Fig. 1(b). We have following assumptions of cell growth and interaction for the reaction-diffusion model. Microcolonies are shaped in circles centered at each well and grow within the well boundary. Cells grown in the same well have identical growth dynamics and function as one agent. AHLs diffuse fast and freely in space as well as across the cell membrane and get degraded at a constant rate, therefore AHL concentration forms a spatial distribution, as shown in Fig. 1(c).

Our proposed model consists of 2*N* sets of ODEs of intracellular chemical reactions and 2 sets of PDEs describing AHL diffusion and degradation in the environment. Species and reactions are shown in Table. I. The model is written as

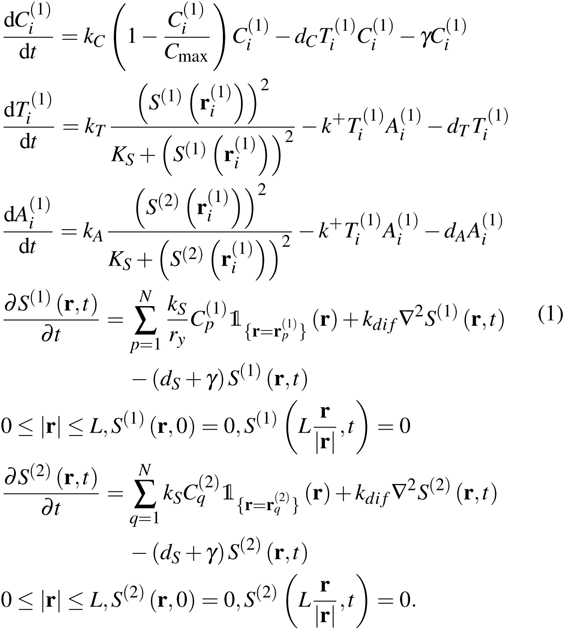

**Table I.**
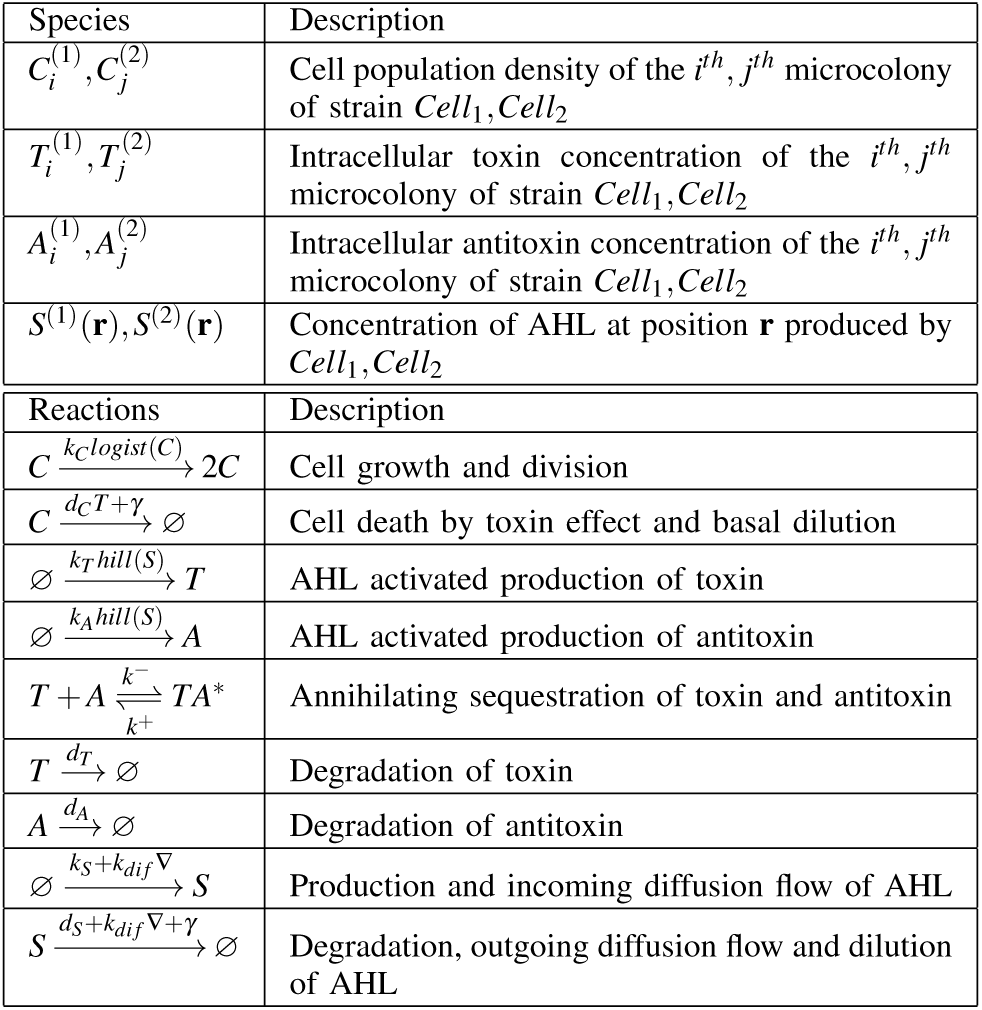
Species and Biochemical Reactions.

Equation (1) describes the dynamics of cell growth and toxin-antitoxin kinetics of *Cell*_1_ in microcolonies indexed by *i* = 1,2,…, *N*, and reaction-diffusion equations and boundary constraints for two AHLs. The production rate of two AHLs are tunable by external IPTG induction, and the induction strength functions as reference to the system. Here, parameter *r*_*y*_ represents the relative induction strength to two cell strains and sets the reference for cell population ratio. Similar dynamics of *Cell*_2_ and intracellur chemical kinetics in microcolonies indexed by *j* = 1,2,…, *N* exist. They are not presented here.

For well-mixed consortia, shown in Fig. 2(a), the reaction-diffusion model reduces to ODEs by setting the distance between wells to be value 0 and having identical dynamics of all cells of the same strain. Fig. 2(b) shows that the steady state of the relative ratio between two populations matches reference with nearly zero error.

**Fig. 2.**
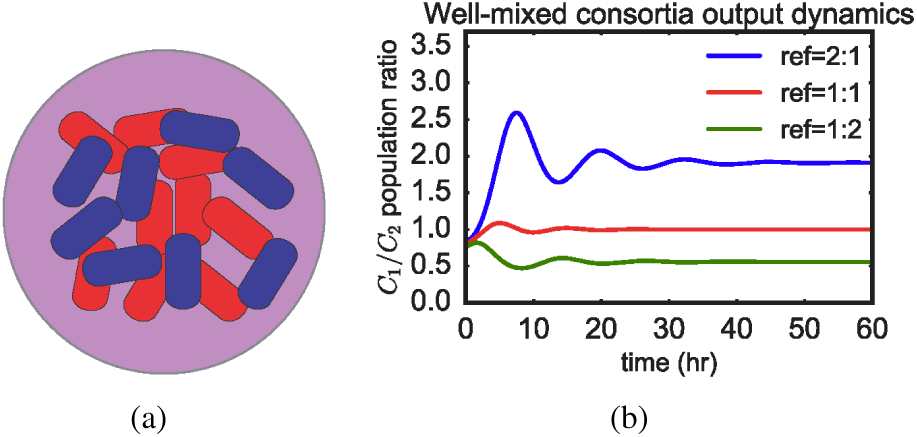
Well-mixed consortia with fractional population control circuit. (a) is a sketch for physically well-mixed consortia. Two cell populations are represented in blue for *Cell*_1_ and red for *Cell*_2_, and the purple background shows the well-mixed environment of two AHLs. (b) describes the output dynamics with different references set by external IPTG inducer levels. The output converges to reference with negligible error at steady state.

For spatially distributed consortia, we can reduce the full model into a simplified one that captures intra/interstrain interactions. We first reduce the PDEs using quasisteady state assumptions, and then reduce the ODEs by linearization. AHL molecules diffuse fast on an agar plate [17] so the timescale of AHL dynamics is smaller than cell growth dynamics. We assume the spatial distribution of AHL reaches a quasi-steady state. Since AHL molecules only affect cell growth when they get absorbed and react with intracellular chemicals, we solve for AHL quasi-steady state at each *Cell*_1_ microcolony center as

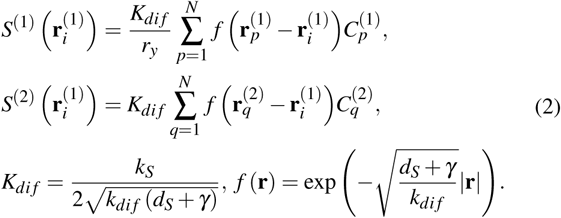

The cell population is regulated via toxin and antitoxin. Since the toxin-antitoxin pair performs strong sequestration, there is a timescale separation of toxin dynamics from cell growth [18]. By performing quasi-steady state approximation and linearization on ODEs in equation (1), we obtain the linearized model in forms of

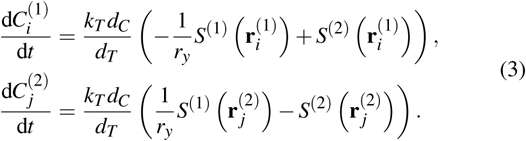

We plug equation (2) into equation (3), and obtain the simplified model as

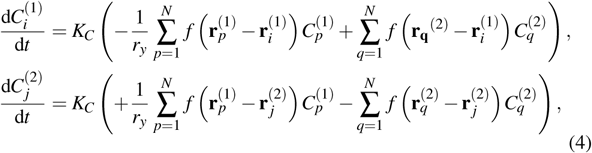

where 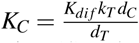.

Equation (4) only contains cell population at discrete well positions as variables. *K*_*C*_ is a lumped parameter characterizing cell growth rate. In both *Cell*_1_ and *Cell*_2_ dynamics, there is a negative term representing intra-strain interaction because cells of the same strain would activate self-killing to avoid growth explosion; there is a positive term representing inter-strain interaction because cells get rescue from the other strain to keep growing. All interaction strengths are functions of the distance between involved microcolonies, so that the model can be derived into a spatial network formula.

## IV. Network Model and Topology Properties

We represent a network model for spatially distributed consortia via a fully connected graph *G* = {*V*, *E*} where *V* = {1,2,· · ·,2*N*} and *E* ∈ *V* × *V* are sets of vertices and edges. The vertices are microcolonies and edges are cell-to-cell interactions via quorum sensing. We define the weighted adjacency matrix *D* = [*d*_*ij*_] ∈ ℝ^2*N*×2*N*^. The element *d*_*ij*_ represents the interaction strength associated with the corresponding edge in *E*.

The network graph can be partitioned into four subnetwork graphs: *G*_11_, *G*_22_ containing microcolonies of one cell strain as nodes and interaction edges among nodes, and *G*_12_, *G*_21_ containing microcolonies of both strains as nodes but only interaction edges between different strains. The matrix *D* can be partitioned into four blocks and they are referred as intra-strain interaction strength matrix –*D*_12_, –*D*_21_, and inter-strain interaction strength matrix *D*_12_, *D*_21_. We associate a set of real values of cell population densities of each microcolony as system states, denoted as 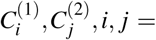, and combine these states into a big network state vector 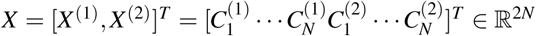. The output *y* is the relative ratio between two cell population densities. We use *l*_2_ norm for all following discussions.

We identify the element di *j* of matrix *D* as *K*_*C*_*f*(**r**_*j*_ – **r**_*i*_) according to equation (4) and derive the network state dynamics as in

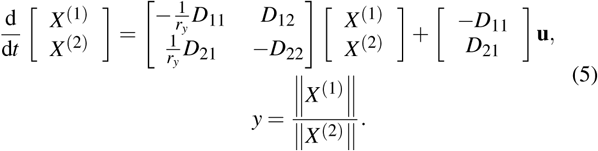

Our control goal is to set the output to track an external reference *r*_*y*_, set by relative IPTG inducer level. We add a network control input 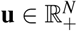 into the system, and it is implemented as an extra production of *AHL*_1_ in each *Cell*_1_ well. Therefore, the coefficient matrix of **u** depends on *D*_11_ amd *D*_21_ because they represent interactions activated by *Cell*_1_. When two cell populations are grown on a plate, the spatially structured interaction network may require a control input **u** to steer the system to the steady state that satisfies the reference *r*_y_. We define the control cost of **u** as 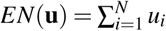, where *u*_*i*_ are elements of **u**, *i* = 1,2,···,*N*. Consider a network *G* = ⋃*G*_*ij*_, *i*, *j* = 1,2 with adjacency 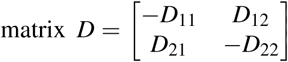. We define properties such as symmetry, locality and connectivity in the context of quorum sensing in microbial consortia.

### Definition 1.

If D_11_ = D_22_, D_12_ = D_21_, then G is a symmetrical network.

*Remark*. Given the assumption that two AHLs diffuse and get degraded at the same rate, the geometric symmetry of spatially distributed consortia is equivalent to network topology symmetry.

### Definition 2.

The locality of a subnetwork G_ii_ is defined as ||D_ii_||, i = 1,2.

*Remark*. Locality measures the intra-strain interaction strength. If microcolonies are grown closer on the plate, they form a more localized community.

### Definition 3.

The connectivity of a subnetwork G_ij_ is defined as ||D_ij_||, i, j = 1,2, i ≠ j. Connectivity of a network G is defined as ||D_ij_ 11 + ||D_ji_||, i, j = 1, 2, i ≠ j.

*Remark*. Connectivity represents the inter-strain interaction strength. If two cell populations are more mixed than isolated, the network is strongly connected with large connectivity. Otherwise, the network is weakly connected.

## V. Relationship Between Topology and System Behaviors

The proposed synthetic circuit ensures good performance in regulating relative ratio of cell population for well-mixed consortia. The results in Fig. 2(b) and from previous work shows that the output *y* matches the reference *r*_*y*_ with negligible error [11]. However, The output’s accurate tracking performance does not maintain for spatially distributed consortia and an extra network control input is needed. By designing the interaction network topology, we can minimize the cost to achieve population control by solving the following problem.

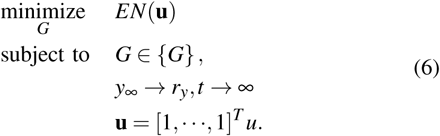

We first set the network control input **u** to be zero and solve for output dynamics of all potential network topologies to see the limits on performing good tracking. Secondly, we look for certain topologies that need small control efforts by solving equation (6). Then we can show how network properties of symmetry, locality and connectivity contribute to the minimum required network control cost.

Given a fixed network topology, there usually exist cost favorable nodes that require less control effort than other nodes to achieve good tracking. It could help us efficiently selecting microcolonies to induce extra AHL production as control input by solving the following equation.

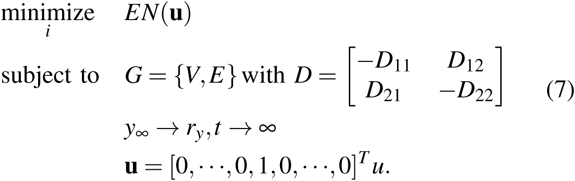

In this section, we present propositions, and give math proof sketch and simulation results. To better demonstrate, we assume the inter-strain interaction strength between adjacent microcolonies from *Cell*_1_ to *Cell*_2_ is equal to the strength from *Cell*_2_ to *Cell*_1_, in other words, 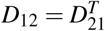; all microcolonies have fixed position within the 2*N* wells, so we have ||*D*|| = *const*; there is an unique steady state solution for all network topologies, denoted as *X*_∞_ and *y*_∞_.

### A. Network symmetry determines system tracking accuracy

#### Proposition 1.

If network G is symmetrical, then the system in equation (5) requires no network control cost to achieve good tracking of reference.

*Proof* Since *G* is symmetrical, we have *D*_11_ = *D*_22_, *D*_12_ = D_21_. Let **u** = **0**, the system dynamics is reduced to

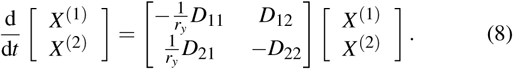

By subtraction of *X*^(1)^ and *X*^(2)^ dynamics in equation (8), we can obtain the dynamics of Δ*X* = *X*^(1)^ – *X*^(2)^ as

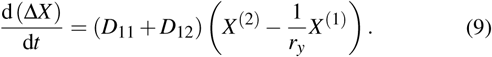

The steady state is solved and it matches the reference.

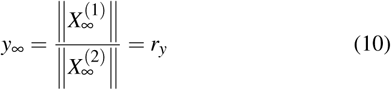

*Simulation* We run simulations on the full reaction-diffusion model in equation (1). We set *N* = 4, *r*_*y*_ = 1 and **u** = **0**. We present three different network topologies that are symmetrical, labeled as sym1, sym2 and sym3 in graph representations and show the system state values (cell population densities on 2D space) at steady state in Fig. 3(a)-3(c). Fig. 4(a) shows the output’s dynamics. Computational results verifies the proposition where the output converges to value 1 without extra network control cost.

**Fig. 3.**
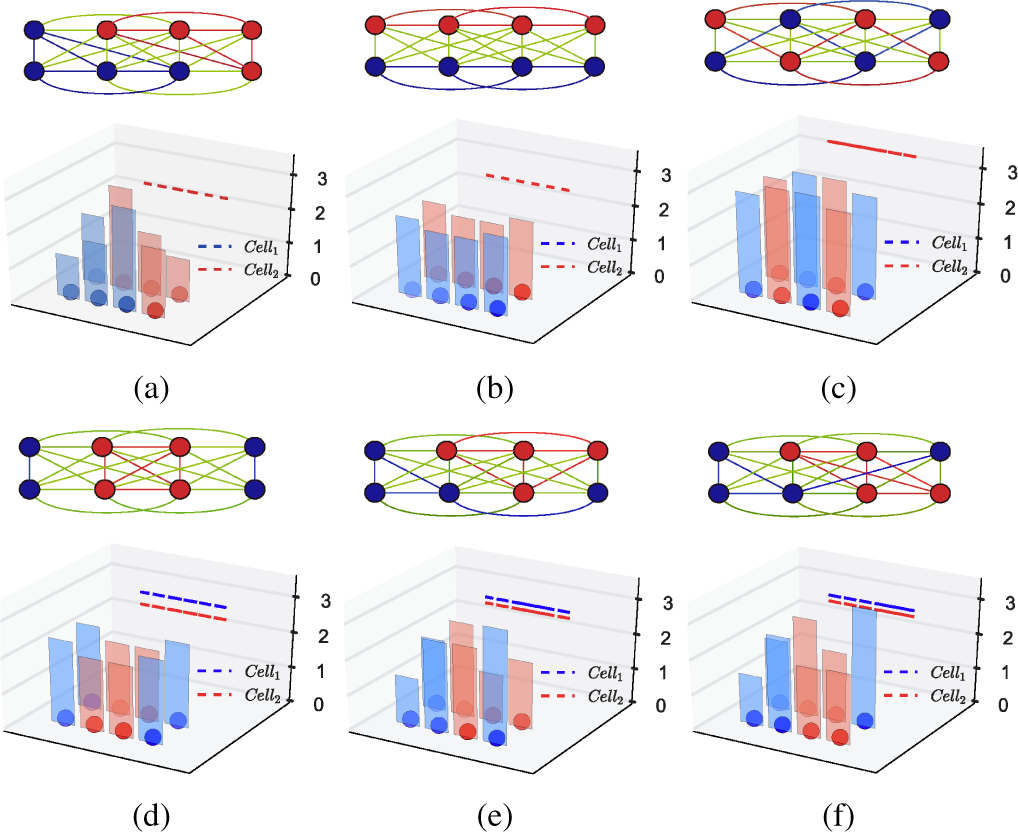
System steady states for spatially cultured consortia with symmetrical or asymmetrical interaction networks. (a)-(c) describe consortia structures with different symmetrical network topologies, denoted as sym1, sym2 and sym3. The upper plots are network graph representations. Blue and red nodes are microcoloinies of two cell populations; blue and red edges are intra-strain interaction pathways; green edges are inter-strain interaction pathways. The lower plots show cell population density at steady state. The bars are population densities of each microcolony and the dashed lines (blue and red) on *x* – *z* plane are total *Cell*_1_ population and total *Cell*_2_ population. The dashed lines overlap, indicating that the relative population ratio between two strains is value 1, so *y*_∞_ = *r*_*y*_. (d)-(f) describe consortia structures with different asymmetrical network topologies, denoted as asym1, asym2 and asym3. The dashed lines are not at same level, indicating that the output doesn’t perform good tracking and the system needs an network control input to achieve referenced cell population fraction.

**Fig. 4.**
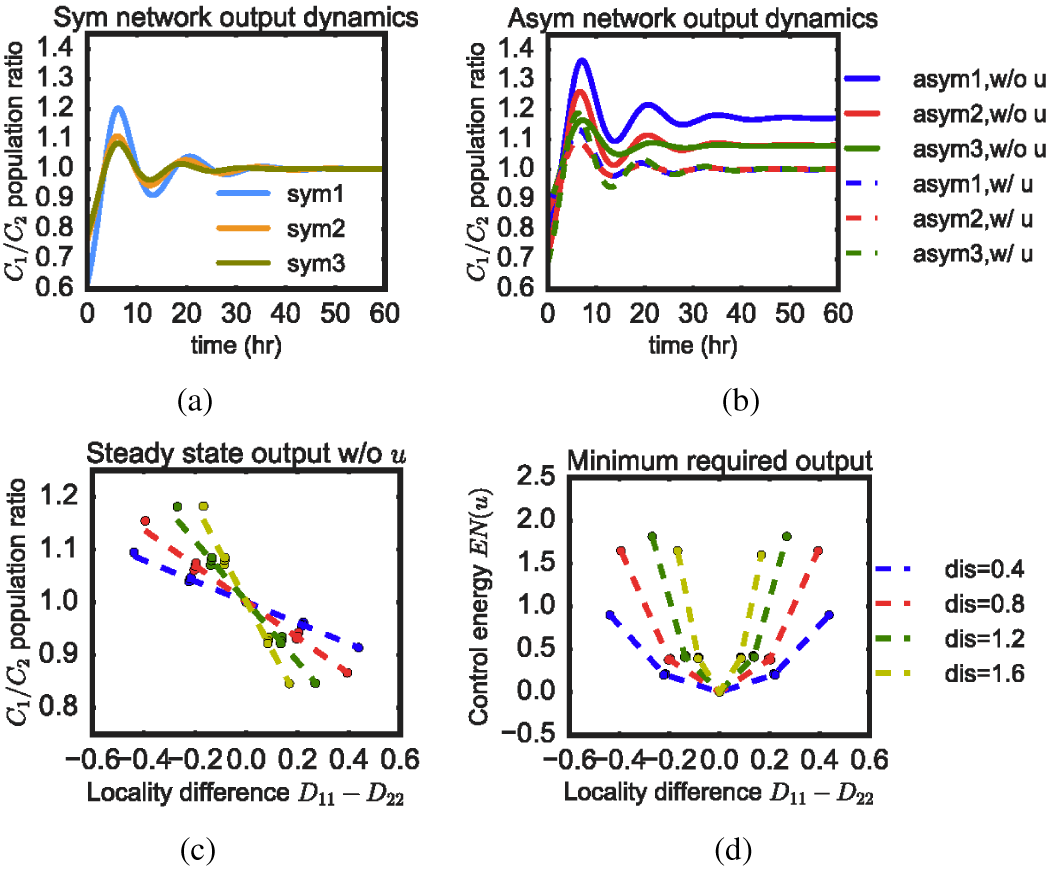
System state and output dynamics w/ and w/o network control input. (a) describes output dynamics of three different symmetrical network topologies (sym1-3) without network control input u. Outputs of all symmetrical networks achieve reference at value 1. (b) describes output dynamics of three different asymmetrical network topologies (asym1-3). Solid lines represent outputs with no network control input *u* and there are significant steady state errors. Dashed lines represent outputs with minimum network control input *u* and all of them converge to the constant reference *r*_*y*_ = 1. (c) describes the relationship between steady state output and the difference between interaction network localities of two cell populations. The dots are simulated data from all potential topologies of consortia cultured in 2*N* = 2 × 4 wells, and the dashed lines are fitted linear functions. Different colors represent results for different physical distance unit values between two adjacent wells. The plot shows that more localized cell population tends to occupy a lower population fraction. (d) describes the relationship between minimum required network control cost and the difference between interaction network localities of two cell populations. The plot shows networks that have large locality difference require more control effort to track reference.

### B. Locality identifies topology that requires minimum control cost

#### Proposition 2.

Given a network G, the minimal network control cost for system in equation (5) is achieved by topologies with minimal locality difference between subnetworks G_11_ and G_22_.

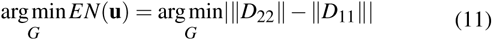

*Proof* Let **u** = [1, ···, 1]^*T*^*u* By subtraction of *X*^(1)^ and *X*^(2)^ dynamics in equation (5), we can obtain

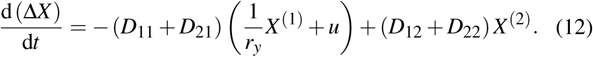

Since there exist a unique solution at steady state, we have *D*_11_ + *D*_21_ is invertible and solve for the steady state.

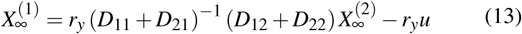

To achieve reference, we let *y*_∞_ = *r*_*y*_ and solve for the minimum required control input cost by giving inequalities.

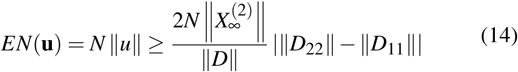

When ||*D*_22_|| – ||*D*_11_|| = 0, one special case is discussed in Proposition 1 and we have *EN*(**u**) = 0 ≥ 0. We can minimized the control cost by minimizing network locality difference.

#### Simulation

We present three different network topologies that are asymmetrical, labeled as asym1, asym2 and asym3 in Fig. 3(d)-3(f). For all these topologies, *Cell*_2_ are more locally distributed than *Cell*_1_, which satisfy ||*D*_22_|| – ||*D*_11_|| > 0. Fig. 3(d)-3(f) demonstrate the system state values (cell population densities on 2D space) at steady state without network control input **u**. They show *Cell*_2_ population occupies a lower fraction than Cell_1_, in other words, *y*_∞_ > *r*_*y*_. We then run simulations with network control input added to the systems and compare the output dynamics in Fig. 4(b). All three systems perform good tracking of the reference with different levels of control cost. We analyze all potential topologies for *N* = 4 spatial settings and plot the relationship between locality difference and steady state output without control input **u** in Fig. 4(c), and the relationship between locality difference and minimum required control cost in Fig. 4(d). We altered the absolute distance unit values between adjacent wells and the relationships maintain.

### C. Locality and connectivity contribution identifies energetically favorable controller node

#### Proposition 3.

Given a network G, the selection of the most cost favorable node depends on the node’s contribution to locality and connectivity.

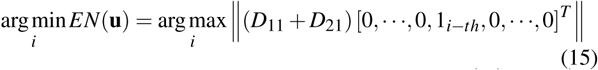

Proof Let **u** =[0,···,0,1_*i*–*th*_,0, ···,0]^T^ *u*. Let 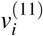 and 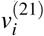 denote the *i*^*th*^ column of *D*_11_ and *D*_21_ as

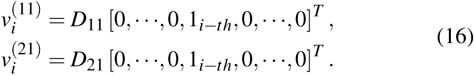

By subtraction of *X*^(1)^ and *X*^(2)^ dynamics in equation (5), we can obtain Δ*X* = *X*^(1)^ – *X*^(2)^ as

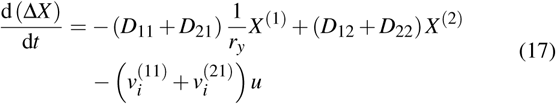

We can solve for the solution as

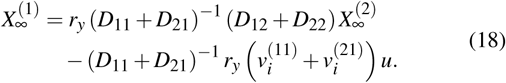

To achieve reference, we let *y*_∞_ = *r*_*y*_ and solve for the minimum required control input cost as

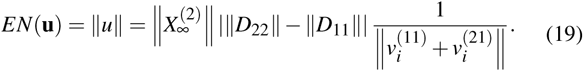

Given a network topology, *D* is fixed and we can minimize the control cost by picking one favorable node that contributes most to locality and connectivity.

#### Simulation

We demonstrate the proposition by adding a constant network control input onto each node of a fixed network topology, and measure the control cost required for steering the system output to track the reference. Fig. 5 shows an example of asym1 network topology. The results show that the maximizer and minimizer of function ||(*D*_11_ + *D*_21_) [0, ···,0,1_*i*–*th*_,0, ···,0]^*T*^|| are the most and least cost favorable nodes. The data show 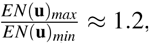, so the worst selection of control nodes can result in 20% more of cost burden.

**Fig. 5.**
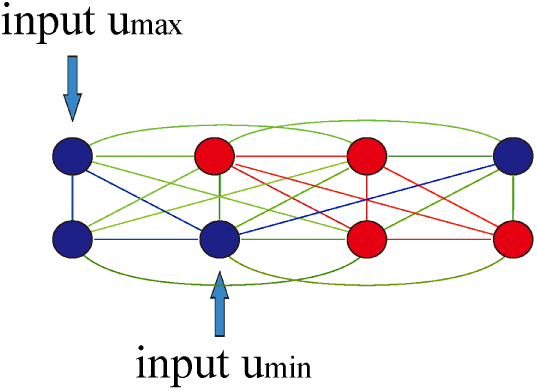
Illustration for picking energetically favorable nodes for asymmetrical networks. The most energetically favorable node is the one in the network center, which contributes most to network’s locality and connectivity. It has a more significant effect on system dynamics because it is strongly connected with other nodes.

### D. Connectivity improves network robustness

In microbial consortia, fluctuations of chemicals in environment can cause uncertainty and noises in cell growth. To investigate the robustness of our synthetic circuit and interaction network, we introduce a metabolic burden change as a disturbance to cell growth, and assess system output’s adaptation as a measure of robustness. For all potential network topologies, we decrease the growth rate of *Cell*_1_ populations after the system achieves some steady states, and measure the output recovery error when the systems achieves steady states again. Fig. 6(a) shows that topologies of large connectivity are more adaptive to perturbations. Fig. 6(b) shows that the most robust network topology is sym3 where all *Cell*_1_ and *Cell*_2_ microcolonies are strongly connected, and the least robust network topology describes consortia that contain two rather isolated populations.

**Fig. 6.**
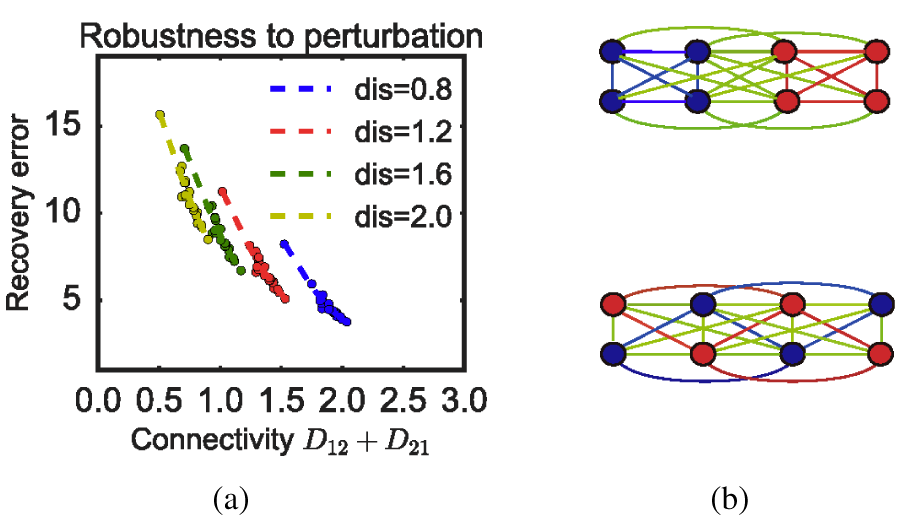
Robustness of networks under perturbations. (a) describes the relationship between network connectivity and output recovery error after *Cell*_1_ growth rate is perturbed to be 20% slower. The dots are simulated data from all potential topologies of consortia cultured in 2*N* = 24 wells, and the dashed lines are fitted linear functions. Different colors represents for different physical distance unit values between two adjacent wells. The plot shows that strongly connected networks are more robust than weakly connected networks with smaller output recovery error. Strongly connected networks have better adaptation to fluctuations in environments. (b) shows two network topologies that have the smallest(upper) and largest(lower) connectivity. Their corresponding systems are the least and most robust one over all potential networks.

From an ecological perspective, our proposed population regulation circuit performs a similar mutualistic symbiosis. Research has shown that mixed consortia are more robust to environmental perturbations and more resistant to invaders [7]. Our model captures such property by relating interaction network connectivity to robustness. In network graph representation, it is clear to see strongly connected topologies correspond to mixed consortia and weakly connected topologies correspond to isolated consortia.

## VI. Discussion

In this paper, we propose a network control strategy to achieve population regulation for spatially distributed microbial consortia. Given an intracellular synthetic circuit, we focus on a physically meaningful understanding of how cell-to-cell interaction network topology governs control potential. We build a network model based on reaction-diffusion equations, map system behaviors to topology properties and formulate the network control question as optimization problems. The propositions inform links from topology to dynamics analytically and numerically, and provide network level design guidance for better performance with less control efforts in synthetic biology problems.

We note that our work is based on a specific synthetic circuit, but the theory and analysis can be adopted to a more generalized problem in synthetic biology. For any functional intracellular circuit that requires cell-to-cell interaction, our approach is useful for finding out constraints on spatial-temporal dynamics. It is important to discuss about such theoretical limitations on population level synthetic system design, such as biofilm formation. For future work, we would like to gain more intuition about the role of topology in complex networks by applying our results to other systems in microbial consortia.

## VII. Acknowledgments

The authors would like to thank Fangzhou Xiao and Leo Green for their insightful discussion. The authors X. R is partially supported by the Air Force Office of Scientific Research, grant number FA9550-14-1-0060. The project depicted is also sponsored by the Defense Advanced Research Projects Agency (Agreement HR0011-17-2-0008). The content of the information does not necessarily reflect the position or the policy of the Government, and no official endorsement should be inferred.

## References

[1] L. Keller and M. G. Surette, “Communication in bacteria: an ecological and evolutionary perspective,” Nature Reviews Microbiology, vol. 4, no. 4, p. 249, 2006.

[2] F. K. Balagaddé, H. Song, J. Ozaki, C. H. Collins, M. Barnet, F. H. Arnold, S. R. Quake, and L. You, “A synthetic escherichia coli predator-prey ecosystem,” Molecular Systems Biology, vol. 4, no. 1, 2008.

[3] W. Weber, M. Daoud-El Baba, and M. Fussenegger, “Synthetic ecosystems based on airborne inter-and intrakingdom communication,” Proceedings of the National Academy of Sciences, vol. 104, no. 25, pp. 10 435–10 440, 2007.

[4] D.-K. Ro, E. M. Paradise, M. Ouellet, K. J. Fisher, K. L. Newman, J. M. Ndungu, K. A. Ho, R. A. Eachus, T. S. Ham, J. Kirby, M. C. Y. Chang, S. T. Withers, Y. Shiba, R. Sarpong, and J. D. Keasling, “Production of the antimalarial drug precursor artemisinic acid in engineered yeast,” Nature, vol. 440, no. 7086, pp. 940–943, 2006.

[5] J. J. Minty, M. E. Singer, S. A. Scholz, C.-H. Bae, J.-H. Ahn, C. E. Foster, J. C. Liao, and X. N. Lin, “Design and characterization of synthetic fungal-bacterial consortia for direct production of isobutanol from cellulosic biomass,” Proceedings of the National Academy of Sciences, vol. 110, no. 36, pp. 14 592–14 597, 2013.

[6] W. C. Fuqua, S. C. Winans, and E. P. Greenberg, “Quorum sensing in bacteria: the luxr-luxi family of cell density-responsive transcriptional regulators.” Journal of bacteriology, vol. 176, no. 2, p. 269, 1994.

[7] A. E. Escalante, M. Rebolleda-Gómez, M. Benítez, and M. Travisano, “Ecological perspectives on synthetic biology: insights from microbial population biology,” Frontiers in microbiology, vol. 6, p. 143, 2015.

[8] M. Newman, Networks: an introduction. Oxford university press, 2010.

[9] Y. Chen, J. K. Kim, A. J. Hirning, K. Josić, and M. R. Bennett, “Emergent genetic oscillations in a synthetic microbial consortium,” Science, vol. 349, no. 6251, pp. 986–989, 2015.

[10] G. Fiore, A. Matyjaszkiewicz, F. Annunziata, C. Grierson, N. J. Savery, L. Marucci, and M. di Bernardo, “Design of a multicellular feedback control strategy in a synthetic bacterial consortium,” in Decision and Control (CDC), 2016 IEEE 55th Conference on. IEEE, 2016, pp. 3338–3343.

[11] X. Ren, A.-A. Baetica, A. Swaminathan, and R. M. Murray, “Population regulation in microbial consortia using dual feedback control,” in Decision and Control (CDC), 2017 IEEE 56th Annual Conference on. IEEE, 2017, pp. 5341–5347.

[12] R. D. McCardell, S. Huang, L. N. Green, and R. M. Murray, “Control of bacterial population density with population feedback and molecular sequestration,” bioRxiv, p. 225045, 2017.

[13] E. M. Bahassi, M. H. O’Dea, N. Allali, J. Messens, M. Gellert, and M. Couturier, “Interactions of ccdb with dna gyrase inactivation of gyra, poisoning of the gyrase-dna complex, and the antidote action of ccda,” Journal of Biological Chemistry, vol. 274, no. 16, pp. 10 936–10 944, 1999.

[14] K. J. Aström and R. M. Murray, Feedback systems: an introduction for scientists and engineers. Princeton university press, 2010.

[15] C. Briat, A. Gupta, and M. Khammash, “Antithetic integral feedback ensures robust perfect adaptation in noisy biomolecular networks,” Cell systems, vol. 2, no. 1, pp. 15–26, 2016.

[16] N. Olsman, A.-A. Baetica, F. Xiao, Y. P. Leong, J. Doyle, and R. Murray, “Hard limits and performance tradeoffs in a class of sequestration feedback systems,” bioRxiv, 2017. [Online]. Available: https://www.biorxiv.org/content/early/2017/11/20/222042

[17] A. W. Decho, R. L. Frey, and J. L. Ferry, “Chemical challenges to bacterial ahl signaling in the environment,” Chemical Reviews, vol. 111, no. 1, pp. 86–99, 2011, pMID: 21142012. [Online]. Available: https://doi.org/10.1021/cr100311q

[18] Y. Qian, T. W. Grunberg, and D. Del Vecchio, “Multi-time-scale biomolecular ‘quasi-integral’controllers for set-point regulation and trajectory tracking,” in 2018 Annual American Control Conference (ACC). IEEE, 2018, pp. 4478–4483.

